# Robust, general purpose, digital power line hum filter which is free of deformations and which can be applied to large transients

**DOI:** 10.1101/830463

**Authors:** Marc H. E. de Lussanet de la Sablonière

**Affiliations:** University of Munster

**Keywords:** biological signals, EEG, EKG, EMG, force plate, hum noise, kinetics, power line interference

## Abstract

Power line interference (“hum noise”) is a common source of noise in recorded biological data. It has a highly constant frequency with harmonics and little variation in amplitude and wave shape. In contrast to stochastic noise it is phasic, i.e., the phase relation remains constant across long intervals. In digital recordings, the measurement frequency is typically very close to a multiple of the hum frequency (50 or 60 Hz). Common filters introduce various kinds of distortions and errors. The here proposed subtraction method makes use of the specific properties of power line hum. For this, it computes a moving estimate of the hum noise by taking the periodic median of the high-pass filtered signal. The resulting periodic median subtraction (PMS) filter reliably removes hum of any harmonic composition, even if the ground frequency is lacking. The filter is completely free of border artifacts. It does not introduce distortions, even if around sharp transients such as in force plate recordings of jumping. The filter is validated on recorded and artificial data. The errors are quantified. The results also show that the errors of hum filters generally increase with the high-frequency content of the data. Thus, the removal of hum from a force plate recording generally gives better results than from an EMG recoding. Compared to other filters, the errors of the PMS filter are generally lower than the best hum filters currently known.

## 1 Introduction

Digital data recordings, such as EEG, EMG, EKG, kinetics (force plates) etc., are all prone to power line noise. Power line noise is a common and old problem and various filters have been made designed for such noise. Even though power line noise is mostly highly regular in frequency, amplitude and wave form, it has proven notoriously difficult to design a filter that is free of artifacts and deformations, which leaves the original signal intact. Moreover many filters focus on a single narrow frequency band, but power line noise typically has strong harmonic frequencies as well.

Hum problems can be solved by avoiding them. If they cannot be avoided, analog or digital filters can be applied. Thanks to modern computational power and digital storage capacity analog filters are not common anymore. There are two dominant approaches to design computational filters, i.e., spectral filters in the frequency domain and subtraction filters in the time domain (for reviews see Baratta et al., 1998; Daniel and Neagu, 2018; Thalkar and Upasani, 2013; BLRI, 2016).

Frequency based filters aim to eliminate the main frequency (and often also the harmonics) from the noisy signal. A serious problem are the distortion artifacts which not only remove the hum noise, but also change the shape of the signal. These are known as Gibb’s rippling and as phase distortions. Gibb’s rippling is especially problematic in the vicinity of transients in the signal, which are especially prominent in force plate measurements (kinetics). Many filters perform badly at the beginning and end of the signal (edge effects), and are thus not well suited for relatively short recordings.

Frequency-based methods involve moving averages technique and IIR notch (kaur Aneja and Singh, 2009), spectrum interpolation (Leske and Dalal, 2019), multi-taper decomposition (Mitra and Bokil, 2007, for multichannel techniques such as EEG) and Fourier decomposition (Singh et al., 2019).

Subtraction-type filters aim to fit the shape and amplitude of the hum noise directly from the data, sometimes from multiple channels (e.g., in EEG signals) or from a reference channel (e.g. Wan et al., 2006; Lin et al., 2016). A major advantage of time-domain filters over frequency-based ones, is the highly regular nature of hum noise. Typically, the recording frequency is a multiple of the power line frequency, so that each period of samples of hum noise is an almost exact repeat of the previous and following periods.

The periodic shape of the hum noise can be estimated using wavelet-independent component analysis (Akwei-Sekyere, 2015; Kaushal et al., 2015; Oliveira et al., 2018), eigenvalue decomposition (Sharma and Pachori, 2018), robust active noise control (Lin et al., 2016), least mean squares (Wan et al., 2006), or recurrent neural networks (Qiu et al., 2017). A very simple but remarkably powerful method is provided by computing the mean period of the hum noise (Levkov et al., 2005).

The goal of the present work is to find a distortion-free hum-filter which is robust to transients in the data and which can handle even short time series. With respect to Levkov et al. (2005), it is substantially improved the method by using a median rather than the mean values for creating the filter, by introducing a moving window. The periodic median is calculated from the high-pass-filtered data and is thus robust to transients.

This new, periodic median subtraction (PMS), filter is tested on artificial and experimental data. As a reference the current best performing frequency-based hum filter is used (Singh et al., 2019).

## 2 Method

Figure 1 illustrates the creation of the filter schematically. The data are high-pass filtered using an 8th-order zero-phase digital Butterworth filter (Matlab functions butter and filtfilt). The cut-off frequency optimally is somewhat below half the power line frequency (i.e., 20 Hz cut-off for 50 Hz power line frequency), to ensure that the high-pass filter does not deform the noise components of the data.

**Fig. 1.**
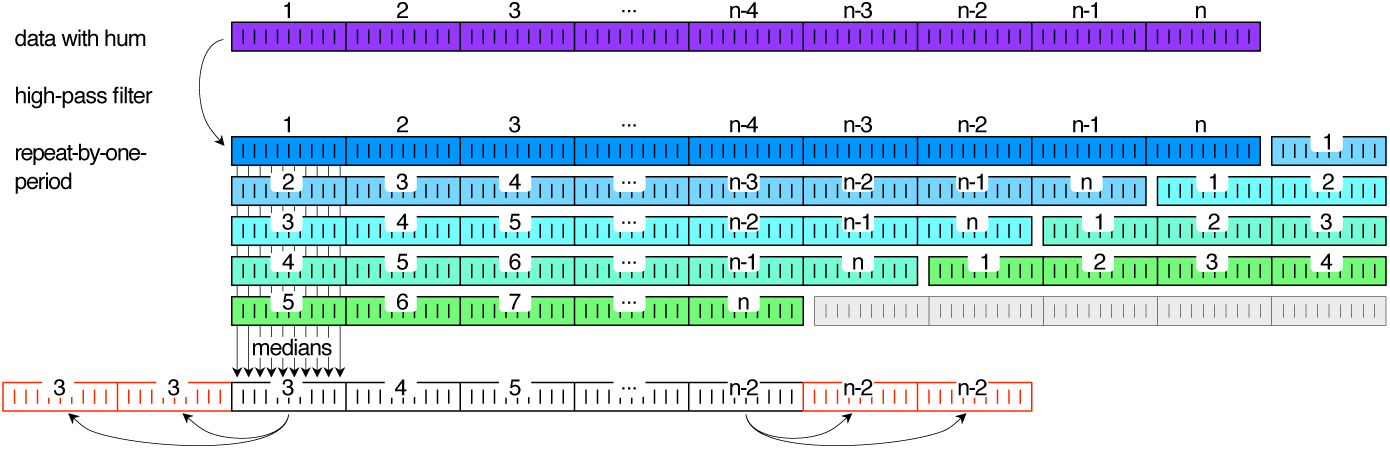
Schema of the periodic median subtraction (PMS) filter. The raw data contain a pattern of additive hum noise of n periods (top row). For construction of the filter, the offset and slower trends are removed by high-pass filtering. The thus filtered signal is repeated into a matrix, of which each row is shifted by one hum period. The number of rows is the width of the filter window, expressed in hum periods. For each column the median is calculated. The first and last periods are repeated at the beginning and end of the filter respectively. The thus obtained filter is subtracted from the data.

Then the periodicity is established as the ratio of the measurement frequency and the power line frequency. If this is not a whole number, the smallest multiple of the period that constitutes a whole number is to be taken).1 The shape of the power line noise for each sample is then estimated as median of the samples differing a multiple of the noise period from the current sample. Recorded signal usually show fluctuations in amplitude (and/or frequency, see footnote 1) and spectral shape. Performance is much improved when the filter is designed as a moving window. The thus retrieved time series (i.e., the filter) is subtracted from the raw data.

We implemented two cases, which are available as a Matlab function (de Lussanet, 2019). If the window of the length of the data (or longer). The array of data is reshaped to a matrix of size *N*_*periods*_ × *N*_*humperiod*_.

For the case of a moving window, an array is created that repeats the data for *n* times, with *n* the width of the window, expressed in periods of the hum. This array is then reshaped into a matrix, which is one period longer than the data. By this reshaping, each consecutive row of data is shifted by one period with respect to the previous one.

The filter is calculated as the median across the columns. The resulting array of the length of the hum period is repeated to be at least as long as the data. Small offsets are corrected for by subtracting the mean.

In the case of a moving window, the beginning and end of the filter matrix overlap beginning and end of the data series. Therefore these parts of the filter are replaced by repeats of the first and last valid period of the filter respectively (Fig. 1).

## 3 Validation

The method was validated using simulated and real data. The simulated data was a sequence of white noise, 100 s, 1000 Hz and an amplitude of 1. In some simulations the simulated data were low-passed filtered at 6 Hz (bidirectional 1st order Butterworth).

The simulated additive noise was a wave-shaped pattern with rich harmonics. It had a period of 20 samples (50 Hz) and an amplitude of 10. In some simulations, other amplitudes (signal-to-noise-ratio, SNR) and other periodic patterns were tested (sinusoidal, periodic random sequence, and periodic pulse).

For testing purpose, a step of 100 was added to the data at 100 s. During the interval 20-22 s, the noise amplitude was reduced from 10 to 8.

The experimental data involved jumping on a force plate and EMG recordings using two different systems. The jump was recorded using Kistler force plates (type 9287CA), connected via a Kistler DAQ (Type 5695A1) to the recording computer. The data wer sampled at a rate of 800 Hz. The DAQ had an external power adapter, whereas the computer had an internal power adapter, so the housing was connected to earthing of the 220 V AC plug. As the housing of the computer and the DAQ were not electrically connected, a considerable 50 Hz hum with some harmonics was present in the recorded data (amplitude *∼*20 N). The recorded drop jump consisted of two loaded phases and two non-loaded phases, with sudden, transient transitions between these phases.

One bipolar EMG recording was made using a wired system (ToM Erfassung, GJB Datentechnik GmbH, Germany) with local amplifiers close to the surface electrodes (disposable Ag/Ag-Cl electrodes type H93SG, Covidien, Neustadt, Germany). The electrodes were applied to the prepared skin over the right *trapezius* neck muscle and the reference electrode was placed over the C7 vertebra. The recording was made in a free-driving vehicle, whereas the power was taken from the on-board battery. The battery’s DC current was transformed to 220 V 50 Hz AC by using a standard DC-AC power transformer. The transformer caused a strong hum in the EMG signal, which was mainly composed of the odd harmonics.

The other EMG recording was made of the right *extensor digitorum longus* muscle, using a wireless bipolar recording (Noraxon TeleMyo DTS) and disposable surface ekectrodes. The recording was made at 1500 Hz with the standard online 500 Hz low-pass filter applied. The signal was completely hum-free. An artificial 50-Hz harmonic-rich hum was added to the recorded data, but various kinds of hum spectra were tested also.

The PMS subtraction filter was applied to the raw data with hum in all cases. The standard window of 50 hum periods (1.0 s) was applied. As a reference, we also applied the FDM filter (the Fourier decomposition method by Singh et al., 2019), omitting 2-Hz windows at all harmonics up to 499 Hz. The EMG data were then rectified, and smoothed using a low-pass 2nd-order Butterworth filter (the order was effectively doubled by applying the filter bidirectionally to prevent phase shifts).

## 4 Results

The results of the simulations are summarized in Figure 2. The original data consisted of white noise with amplitude 1, the filter was applied to these data with added hum with an amplitude of 10 times the signal. The absolute error in terms of the amplitude of the white noise “signal” was 0.136/0.115/0.892/0.103 (mean/median/max/st.dev.) for the new PMS filter. As a comparison, the “gold standard”, the FDM filter (Singh et al., 2019) gave 0.158/0.132/1.382/0.120, which is worse on each of these measures. Panel C shows how well the filtered signal (red curve) represents the original data (green).

**Fig. 2.**
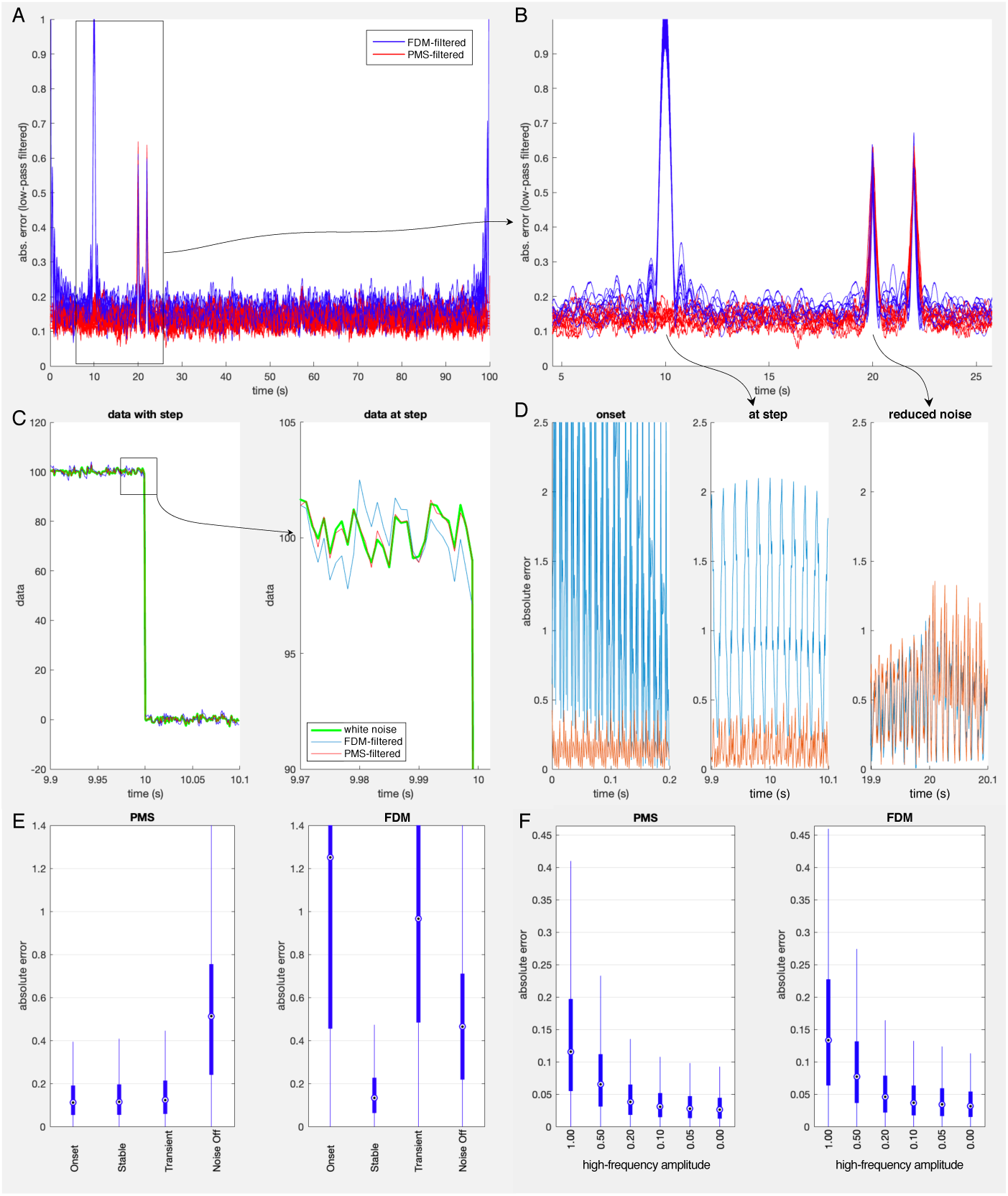
Comparison of the FDM (Singh et al., 2019) and the new PMS hum-filter for simulated data (1000 Hz white noise of amplitude 1, with hum of amplitude 10). At 10 s, there was a step in the data; at 20-22s the noise amplitude was reduced to 8. A. Absolute, smoothed error of the filtered signal with respect to the hum-free data, in 10 simulations. B. Selection of panel A. C. Data without hum and two filtered versions around the step in the data at 10 s of one exemplary simulation and a selection just before the step. D. Absolute (non-smoothed) error at onset, at the step in the data and at onset of reduced noise, for one exemplary simulation. E. Box plots of the absolute error for four time windows (100 simulations). The stable period was taken from 25-99 s, the other windows lasted 0.2 s each (left: the new PMS filter, right the reference FDM filter). F. Median absolute error as a function of the amplitude of high frequencies in the data.

Panel A of Figure 2 shows the smoothed absolute error for ten simulations in both filters. It illustrates how the PMS filter is free of onset deformations and has a very low error for the stationary period. The enlargement in panel B shows the influence of the step in the data and the sudden change in noise amplitude. Note, that the change of noise amplitude (from 10 to 8) was quite large considering that the signal had an amplitude of 1.

In the onset and offset period the performance of the PMS filter was completely stable, as shown by panels D and E (Fig. 2). Also the large, transient step in the data did not affect the filter results (Panel C-E). Only the sudden, large change in hum amplitude caused a considerable error shortly before and after the onset and offset.

The quality of the hum-filtering depended strongly on the amplitude of high frequencies (HF) in the data (Fig. 2F). Applying the PMS hum filter to white noise data (HF amplitude of 1: panels A-E) led to considerable errors (0.115 of the signal amplitude), whereas the errors were small (0.031) for HF amplitudes of 0.1 and below. The same trend was found for the FDM filter (Fig. 2F).

One manner to reduce the PMS filter-errors is to increase the window. Increasing the filter window of the PMS filter from 50 to 500 periods (10 s) reduced the median absolute error in white noise data to 0.037, which is equivalent to the error in a signal with an HF amplitude of 0.2 (cf. Fig. 2F).

Changes of amplitude and shape of the hum-noise gave highly comparable results.

In the example of Figure 3A-F, a power inverter was the source of the hum-noise in an EMG recording. Consequently, the hum consisted almost entirely of harmonics (panel C). The PMS filter completely eliminated the lower harmonics and most of harmonics above 700 Hz (Panel C). Panel B shows a sort period without muscle activity. Panel D shows somewhat more than six periods of the filter. It can be seen how the filter changes slightly and gradually. Panels E and F show the cleaned, rectified EMG before and after low-pass filtering, showing a clean signal with a good resting amplitude, despite considerable hum-noise in the original recording.

**Fig. 3.**
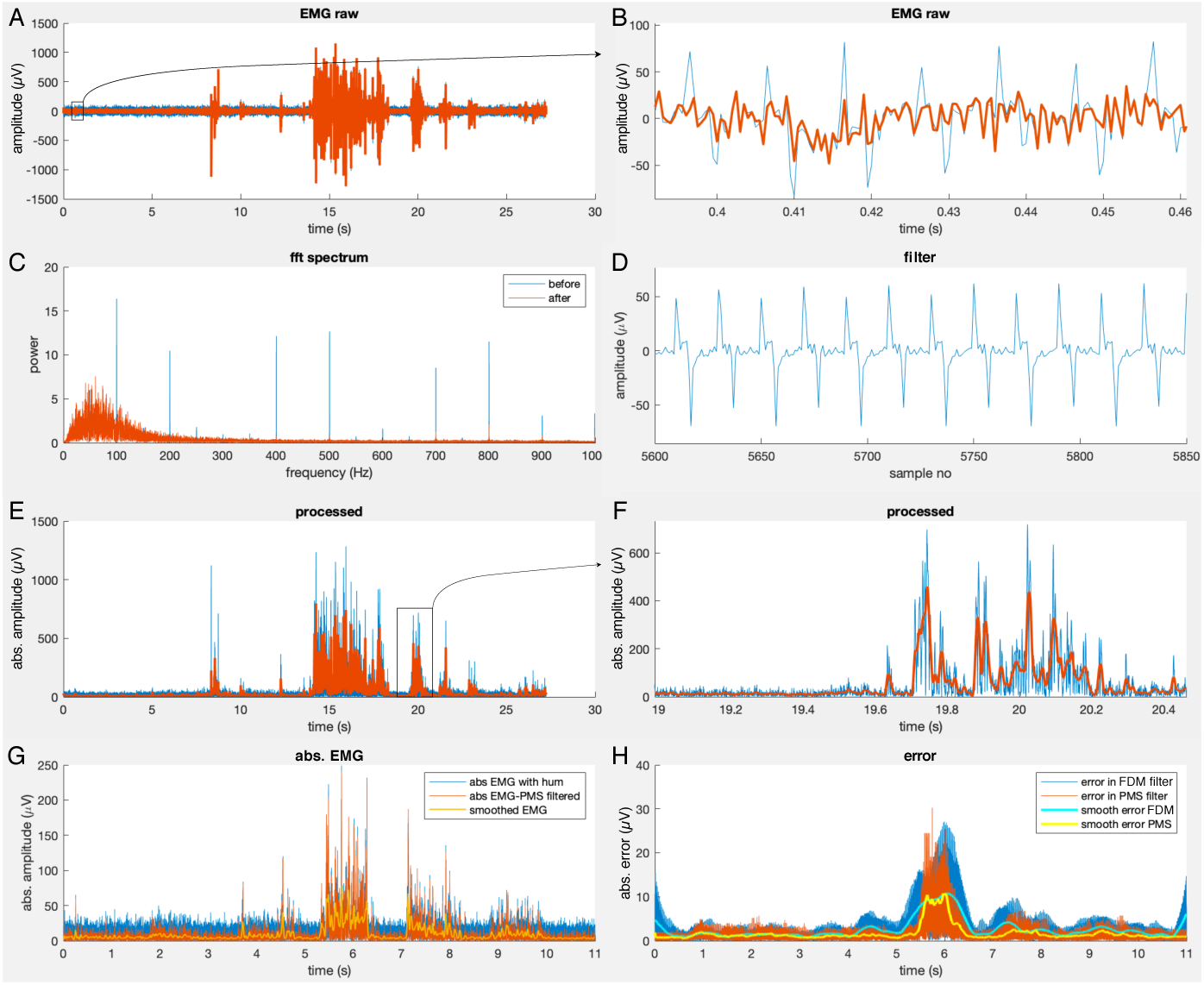
Example of EMG data from the right trapezius muscle, which are contaminated by a DC-AC power inverter. A. Raw data (aligned by a 5 Hz 2nd order butterworth high-pass filter). The peak at 8 s is a movement artifact. B. Selection of A: signal with (blue) and without (red) hum. C. FFT power spectrum of the signals in A. For most frequencies the curves overlap exactly, so the signal with hum (blue) is invisible. D. A short sequence of the extracted hum. Note, that the 50 Hz component is only very minor. E. The signal of A: The hum-cleaned signal rectified (blue) and low-pass filtered at 60 Hz (4th order Butterworth). F. Selection of E. G. EMG of ext. digit. long. with added artificial hum with rich harmonics. The subject subsequently lifted the five fingers beginning with the thumb. H. Absolute error of the FDM filter and the new PMS filter for the EMG recording with hum of panel G.

Figure 3G-H show an EMG recording made with wireless electrodes, which are completely free of power line hum noise. Artificial hum was added to the signal and removed using the new PMS filter. The absolute error of the PMS filter is shown in panel H (red). The yellow trace shows the smoothed error. As a reference, the error of the FDM filter (Singh et al., 2019) is shown in blue. A comparison between panels G and H reveals a correlation between the amplitude of the signal and the error of the filter. For the new PMS filter, the relative error was indeed relatively constant 0.22 / 0.21 (mean / median). This is not as good as for the simulated signal (cf. Fig. 2E, but better than for the FDM filter (0.37 / 0.31 [mean / median]).

In the jumping example (Fig. 4), the amplitude of the hum-noise was as much as 10 N, because the AD converter (Kistler DAQ) was not grounded. In the presented drop jump, the subject landed at first mainly on plate 6 and landed after the jump mainly on plate 3. However in each landing part of one foot contacted the other plate, so both plates registered each landing. In the presence of the hum noise, it would not be possible to determine the exact times of contact and release, but after subtraction of the hum noise, the transients are so sharp that the precise time sample can be determined in each case (panels B and F).

**Fig. 4.**
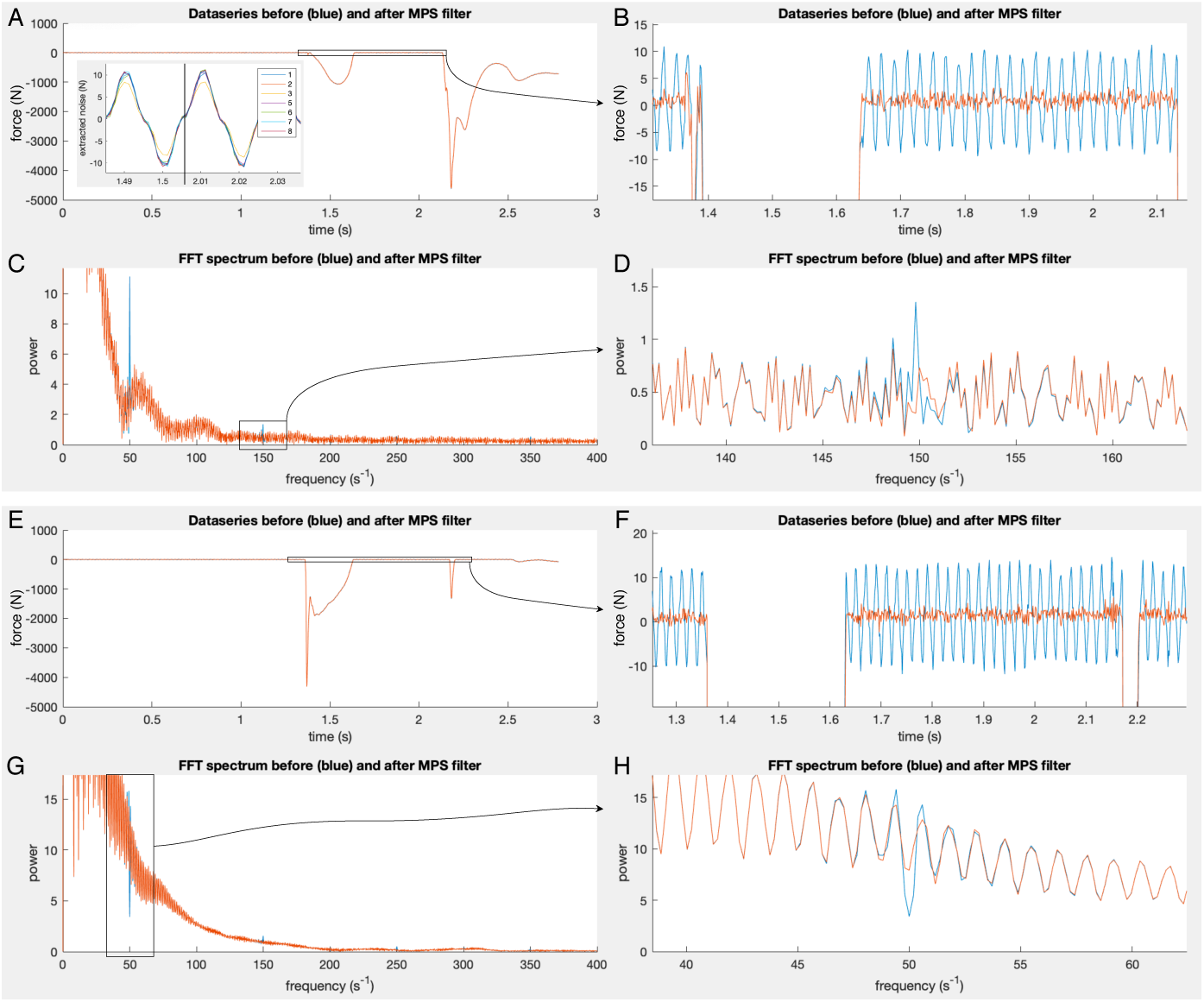
Two examples of the PMS filter, applied to measured vertical force data of a drop jump second (A-D) and first (E-H) landing. Right panels show the selections of the left panels respectively. Shown are the time series of the vertical force and the frequency spectrum of the fast Fourier transformed signal (FFT). The original signal (blue) is affected by substantial power line hum with harmonics, whereas the red curves represent the filtered signal.

The shape of the frequency spectrum of the filtered signal is only changed at the harmonics (panel A, C). At 150 Hz (C) and 50 Hz (D) it can be seen that the frequency is not merely depressed or interpolated. Rather, the pattern of the flanking spectrum is restored in a realistic pattern. This is especially clear in panel D, where the filtered spectrum regularly continues the oscillating pattern in the 50 Hz range. Interestingly, the frequency power at 50 Hz is even enhanced by the filter.

## 5 Discussion

This study presents and validates a new filter for power line hum noise. Power line hum still presents a serious problem in many kinds of measurements, ranging from electrophysiological to force plate recordings. The new method introduces periodic median subtraction (PMS) and presents a number of improvements with respect to existing subtraction methods. The validation showed that the filter also performs better than frequency spectrum-based filter, i.e., the current “gold standard”, the FDM filter (Singh et al., 2019).

An advantage of the PMS filter is that it is completely independent of the harmonic structure of the hum noise. Moreover, by introducing a filter window it is highly tolerant to temporal variations of the shape and amplitude of the hum.

It is not common to analyse the true errors that are caused by filtering the data. Instead, studies tend to plot data before and after filtering in separate plots. The present study chose to rather analyse the true errors made by the filter, and so to quantify the distortions introduced by filtering. An important result is that the high-frequency content of the signal themselves is a major factor in the quality of the filtered result. This was irrespective of the filter used. Thus, for both the new PMS subtraction method, as well as for the reference Fourier method (FDM) the presence of high-amplitude, high-frequency content in the signal resulted in substantially larger errors in the filtered result. This was so for the simulated data as well as for the measured data. Thus, for EMG (and thus likewise for other electrophysiological data) power line hum filtering causes relative large distortions to the data, whereas the same filter applied to force plate measurements is virtually distortion-free.

The new PMS filter performed better than the reference filter on all measures. Theoretically, the performance can even be further improved by increasing the filter window. On the simulated data, the error made to a signal consisting of white noise (cf. Fig. 2F) could be compensated by a ten-fold increase of the window (i.e., from 1 to 10 seconds). In practice however, the hum is usually not so stable that such a long window of 10 s is practicable. In the various data to which the PMS filter was applied so-far, a window of 50 periods (1 s) usually was the optimal compromise (for the force plate data, longer windows seem to be even better, but the difference was so small, that it was irrelevant for the usual applications). With the sudden, large change on noise amplitude, both filters showed considerable distortions.

In the onset and offset as well as in the transient condition the new PMS filter had no increase of the error at all. This is quite unusual. Even though the FDM filter that was used as a reference is robust to small offsets and baseline oscillations, it did show considerable distortion of the signal at onset and offset as well as during the transient. The distortion-free behaviour of the PMS reveal its extraordinary robustness.

### Conclusions

The here-proposed PMS filter provides a simple but very robust method. The filter is free of phase distortions has excellent handling of transients in the data. It automatically handles any harmonic composition of the hum noise. It performs especially well with high amplitudes of hum noise and copes well with variability of phase and amplitude of the noise. Compared to existing subtraction filters, the PMS filter excels in robustness, by using a window, by using the median period for extracting the current hum period, and by being based on the high-passed data. Compared to the Fourier-based filter that was used as a reference (Singh, 2018), the PMS has considerably better properties in all mentioned aspects (onset/offset, transients, noise amplitude variability, low SNR). The PMS also performed better than the FDM on the analysed force and EMG recordings. Concluding, the PMS is probably the best hum filter. A Matlab routine with example data is provided.

## Conflict of interest

The author declares that he has no conflict of interest.

## Open Practices Statement

The data and materials for all simulations are available at (https://www.github.com/lussanet/PeriodicMeanSubtractionForHumNoise). No Experiment was preregistered.

## Footnotes

1 Note, that a small deviation of the frequency, e.g., due to the usage of a power generator or power inverter is not problematic.

## Notes

https://github.com/lussanet/PeriodicMeanSubtractionForHumNoise

